# Protein *N*-glycosylation is essential for SARS-CoV-2 infection

**DOI:** 10.1101/2021.02.05.429940

**Authors:** Aitor Casas-Sanchez, Alessandra Romero-Ramirez, Eleanor Hargreaves, Cameron C. Ellis, Brian I. Grajeda, Igor Estevao, Edward I. Patterson, Grant L. Hughes, Igor C. Almeida, Tobias Zech, Álvaro Acosta-Serrano

**Affiliations:** Department of Vector Biology, Liverpool School of Tropical Medicine, UK; Department of Tropical Disease Biology, Liverpool School of Tropical Medicine, UK; Department of Molecular and Cellular Physiology, University of Liverpool, UK; Department of Biological Sciences, Border Biomedical Research Center, University of Texas at El Paso, TX, USA; Department of Biological Sciences, Brock University, Canada

**Keywords:** SARS-CoV-2, COVID-19, coronavirus, *N*-glycosylation inhibition, drug repurposing, viral infection

## Abstract

SARS-CoV-2 extensively *N*-glycosylates its spike proteins, which are necessary for host cell invasion and the target of both vaccines and immunotherapies. These sugars are predicted to help mediate spike binding to the host receptor by stabilizing its ‘open’ conformation and evading host immunity. Here, we investigated both the essentiality of the host *N*-glycosylation pathway and SARS-CoV-2 *N*-glycans for infection. Inhibition of host *N*-glycosylation using RNAi or FDA-approved drugs reduced virus infectivity, including that of several variants. Under these conditions, cells produced less virions and some completely lost their infectivity. Furthermore, partial deglycosylation of intact virions showed that surface-exposed *N*-glycans are critical for cell invasion. Altogether, spike *N*-glycosylation is a targetable pathway with clinical potential for treatment or prevention of COVID-19.

---

The severe acute respiratory syndrome coronavirus-2 (SARS-CoV-2) causing the coronavirus disease-19 (COVID-19) pandemic is a positive single-stranded RNA enveloped betacoronavirus(*1*). Its homotrimeric spike glycoprotein has been extensively studied as it is the target of most vaccines and treatments under development due to its surface exposure, immunogenicity and essential role in infection. It is well established that the main route for SARS-CoV-2 to infect the host cell is via attachment to the host receptor angiotensin-converting enzyme 2 (ACE-2) through the spike’s receptor-binding domain (RBD), followed by membrane fusion and invasion(*2, 3*). Both spike and ACE-2 are modified by the addition of a variable number of *N*-glycans. While ACE-2 contains seven occupied *N*-glycosylation sites(*4*), the spike protein is heavily *N*-glycosylated with up to 66 oligosaccharides (22 per monomer)(*5, 6*), including vaccine-derived spikes(*7, 8*). Molecular dynamics simulations predict critical roles for spike *N*-glycans in stabilizing the trimer and the RBD open conformation(*5, 6, 9-11*). In addition, they are suggested to be important for binding to ACE-2(*12, 13*) and mutations in specific *N*-glycosylation sites (i.e. N331, N343) drastically reduced pseudotyped virus invasion(*14*). Moreover, spike *N*-glycans are flexible molecules that can occupy a large surface, hiding epitopes from antibodies to escape the host immune response(*14–16*). Considering the important functions *N*-glycans may play, not only from spike but likely from other viral and host proteins, *N*-glycosylation serves as an attractive target to develop new approaches against COVID-19. However, little is known about the functional role of *N*-glycosylation during a real SARS-CoV-2 infection. A few studies have shown the efficacy at reducing viral infection using inhibitors of the endoplasmic reticulum (ER) α-glucosidases(*17-19*) and the Golgi α-mannosidase I(*20*), which are essential enzymes for the maturation of *N*-glycan structures. In addition, knocking out *N*-acetylglucosaminyl-transferase MGAT1 decreased pseudovirus infection(*20*) or resulted in reduced spike binding(*12*). Most of these findings are based on work using heterologous spike expression, pseudotyped vesicular stomatitis virus(*21*) and computational models, but little is known from experimental evidence using actual SARS-CoV-2 virions. In this study, we investigated the essentiality of host *N*-glycosylation and SARS-CoV-2 *N*-glycans for infection *in vitro*.

Considering the potential important roles of spike *N*-glycans, we hypothesized that the partial or complete inhibition of the host *N*-glycosylation pathway may interfere with the normal development of SARS-CoV-2 infection (**Fig. 1A**). This may either prevent the invasion of cells with glycosylation defects in key surface receptors, affect the normal production of viral proteins and virions, and/or hamper their capacity to infect further host cells. In both Vero E6 (African green monkey kidney cells) and HEK293^ACE-2^ cells (lentiviral overexpression of ACE-2 in human embryonic kidney cells), we used RNAi to knockdown the following key enzymes in the *N*-glycosylation pathway prior to SARS-CoV-2 infection: 1) oligosaccharyltransferase’s catalytic subunit STT3 (isoforms A, B and both) to abolish transfer of *N*-glycan precursors to proteins in the ER; 2) GANAB catalytic subunit of the ER α-glucosidase II to prevent the deglucosydation of immature *N*-glycans attached to proteins; and 3) Golgi MGAT1 to block the formation of hybrid and complex *N*-glycans (**Fig. 1A**). As measured by quantification of anti-spike positive cells (**Fig. 1, B to D**), knockdown of both *STT3* isoforms (*siSTT3A+B*) led to an average infection reduction of 55.4% in Vero and 54.2% in HEK cells. Specific knockdowns were then performed separately to determine the contribution of each STT3 isoform. While *siSTT3-A* showed a profound inhibitory effect in HEK cells (52% reduction), it did not significantly reduce viral infection in Vero cells. On the contrary, si*STT3-B* reduced the infection in Vero cells down to 48% but did not protect HEK cells from infection due to inefficient knockdown. Reduction of *GANAB* consistently resulted in less infected Vero and HEK cells, although the effect was not as pronounced as seen in *siSTT3* (41.4% and 42.5%, respectively). Lastly, while *siMGAT1* had no effect in Vero cells, it showed some protection in HEK cells (58.8% infection). We further validated these results by quantifying the amount of infectious viral particles released by Vero siRNA cells (**Fig. 1E**). Knockdown levels were determined for each group and biological replicate by immunoblotting using specific antibodies (**Fig. S1**), which showed that only HEK cells seemed resistant to *STT3-B* knockdown. HILIC-UPLC *N*-glycoprofiling confirmed the expected overall glycan downregulation in si*STT3* and si*GANAB* cells, as well as the accumulation of M5 (Man_5_GlcNAc_2_) glycan in *siMGAT1* cells (**Fig S3B and S4B; Data S1**).

**Fig. 1.**
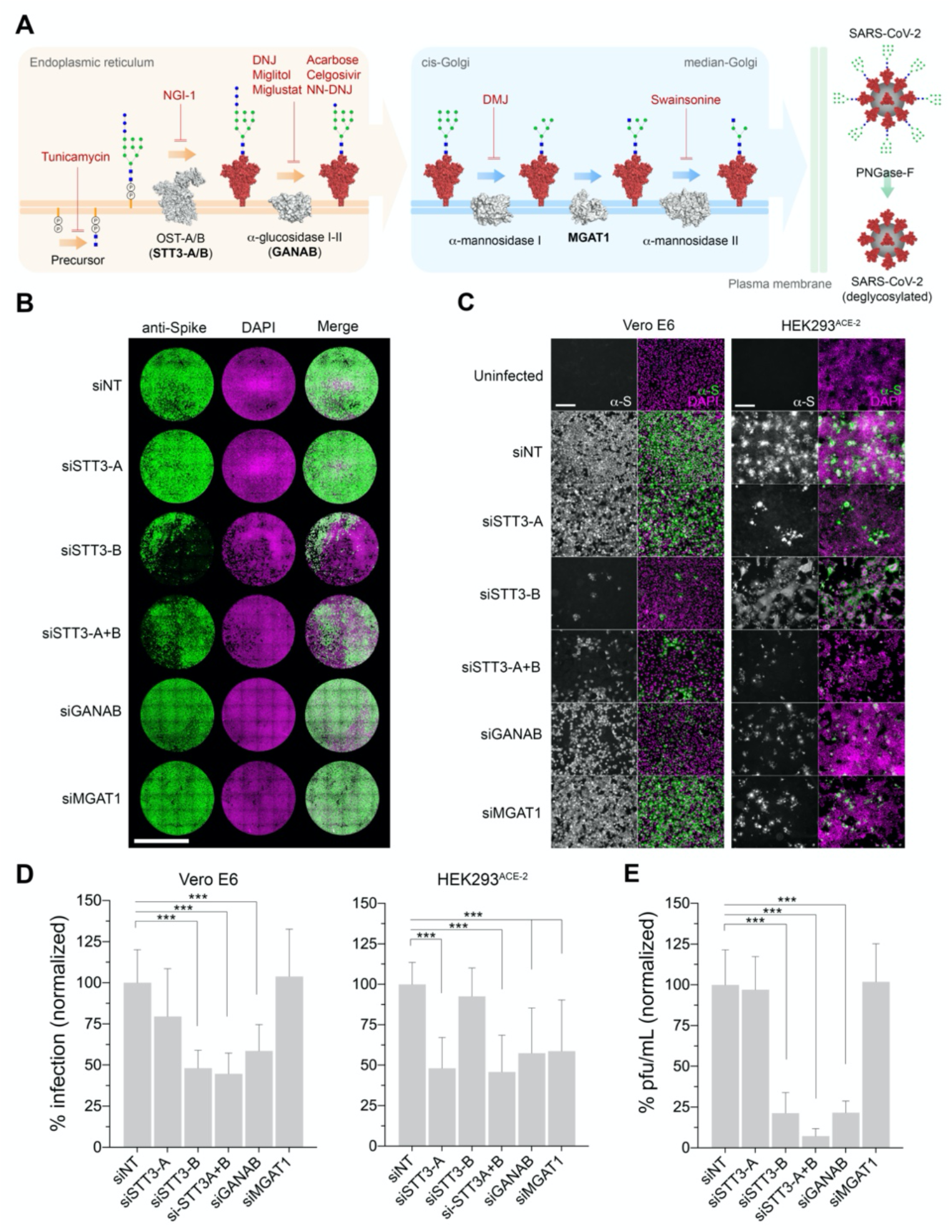
Genetic ablation of host *N*-glycosylation reduces SARS-CoV-2 infection. (**A**) Schematic of the key steps in the *N*-glycosylation pathway targeted in this study. The precursor *N*-glycan is first synthesized in the ER and transferred to the SARS-CoV-2 spike protein (red) by the OST’s catalytic subunit STT3 (isoforms A and B). The *N*-glycans are further processed by the α-glucosidases I and II before the glycoprotein is exported to the Golgi apparatus where the glycans are modified by the α-mannosidase I, MAGT1 and α-mannosidase II. Mature glycosylated virions egress to the extracellular space, where they can be artificially deglycosylated using PNGase F. Glycosylation enzymes (gray); glycosylation inhibitors (red text); glycosylation enzymes targeted by siRNA (bold text). *N*-glycans represented using CFG nomenclature: blue square (N-acetylglucosamine), green circle (mannose), blue circle (glucose); virion *N*-glycan structures are representative only. (**B**) Representative whole-well scans of confluent Vero E6 monolayers transfected either with siNT (non-targeting control) or siRNAs targeting STT3-A, STT3-B, STT3A+B, GANAB and MGAT1, and infected with SARS-CoV-2 (MOI=0.05, 24 hours). Cells immunostained using anti-spike antibodies (green), counterstained with DAPI (magenta) and merged. Scale bar 5 mm. (**C**) Representative images of Vero E6 and HEK293^ACE-2^ siRNA cells infected with SARS-CoV-2 (MOI=0.05 for 24 hours, Vero; MOI=0.1 for 24 hours, HEK293); anti-spike (grey) and merged (green) with DAPI counterstain (magenta). Scale bars 200 μm. (**D**) Percentage of Vero E6 and HEK293^ACE-2^ infected cells (anti-spike positive) in ‘A’; normalized to siNT; three technical replicates, four biological replicates. (**E**) Infectious viral particles in supernatants from infected Vero E6 siRNA cells in ‘A’; percentages normalized to siNT. Bars indicate mean values, error bars represent +S.D., asterisks indicate significance (*p*<0.001).

Having identified several enzymes whose reduction in expression lowered SARS-CoV-2 infections, we then investigated whether chemical inhibition of these and other enzymes throughout the pathway may also promote protection against infection. We tested several compounds known to inhibit important *N*-glycosylation steps, at concentrations considered to be at a low or high end of the effective range (**Fig. 1A**). To emulate the effects of *siSTT3* knocking down the early pathway, we used tunicamycin –to block formation of the *N*-glycan intermediate GlcNAc_2_-dolichol phosphate- and NGI-1 that inhibits STT3 directly. Further down in the pathway as in *siGANAB*, we inhibited the ER α-glucosidase I and II using iminosugars 1-deoxynojirimycin (DNJ), Miglitol(*22*), N-butyl-deoxynojirimycin (Miglustat(*23*)), N-Nonyldeoxynojirimycin (NN-DNJ), Celgosivir(*24*) and Acarbose(*25*). Late in the pathway and resembling *siMGAT1*, we tested the efficacy of Deoxymannojirimycin (DMJ) and Swainsonine to inhibit the Golgi α-mannosidases I and II, respectively. Pre-incubation of both Vero and HEK cells with most inhibitors prior to SARS-CoV-2 inoculation resulted in a significant reduction in infection compared to untreated cells (**Fig. 2, A to C**). Importantly, NGI-1, Miglustat and Celgosivir were equally effective at reducing the infection in either Vero and HEK cells by SARS-CoV-2 variants B.1.1.7 (UK), B.1.351 (South Africa) and P.1 (Brazil) (**Fig. 2D; Fig. S2A**). The same protective effects were observed when the inhibitors were added alongside the virus and up to 4 hours post-inoculation (**Fig. 2E**), and regardless of the inoculum (**Fig. S2, B and C**). For selected inhibitors, we quantified infectious viral titres in cell supernatants (**Fig. 2F**) and measured the proportion of cells showing cytopathic effects (CPE) as a result of the infection (**Fig. 2G**). While many untreated cells became infected and arranged in clusters, both tunicamycin and NGI-1 provided the highest protection rates against SARS-CoV-2 in Vero (97.3% and 97%) and HEK cells (92.3% and 92%) (**Fig. 2, A to C**), showing only few isolated infected cells (**Fig. 2, B and H**). They also protected cells against CPE (94% and 99.5%) (**Fig. 2G**) and at high doses led to undetectable levels of infectious particles in cell supernatants (**Fig. 2F**). Inhibitors of ER α-glucosidases proved to be less effective and presented a greater variability. The most effective ones (i.e., Miglustat, Celgosivir and NN-DNJ) showed reduced overall infections in Vero and HEK cells (**Fig. 2, A to C**), smaller clusters of infected cells (**Fig. 2H**), CPE protection and moderate virus titre reduction in supernatants (**Fig. 2, F and G**). DNJ, Miglitol and Acarbose only showed little effect at the highest doses. While the protection given by tunicamycin and NGI-1 can be attributed to the absence of *N*-glycans, the effects produced by α-glucosidase inhibitors could result from alterations in glycan structure. Alternatively, increased levels of anti-spike in drug-treated cells (**Fig. 2I**) suggests a possible accumulation of viral glycoproteins in the ER as a consequence of the inhibition of the calnexin/calreticulin cycle(*26*). Inhibition of the Golgi α-mannosidases using DMJ and Swainsonine produced no effect in Vero cells but did reduce HEK cell infections, resembling the phenotypes seen in *siMGAT1* (**Fig. 1**). Differences in baseline *N*-glycosylation profiles between Vero and HEK cells (**Fig S4A; File S1 and S2**) and expression of the Golgi’s endo-α-1,2-mannosidase (MANEA)(*27*) independent pathway may account for such variations. Cell viability assays were run in parallel to rule out poor metabolism or toxicity as the main cause of infection decrease (**Fig. S2D**), although drug off-target effects may play a role at high doses. Lastly, we investigated whether two inhibitors combined may increase their potency (**Fig. S2E**). While the combination of tunicamycin and NGI-1 did not significantly improve their respective individual efficacies, simultaneous treatment of NGI-1 with any α-glucosidase inhibitor promoted higher protection rates compared to NGI-1 or α-glucosidase alone. Furthermore, some combinations of α-glucosidase inhibitors were particularly efficacious (e.g., Miglustat combined with Miglitol) and combinations with α-mannosidase inhibitors also resulted in augmented efficacies. The corresponding changes in protein glycosylation upon drug treatment were determined by HILIC-UPLC (**Fig S3A and S4C; Data S1**).

**Fig. 2.**
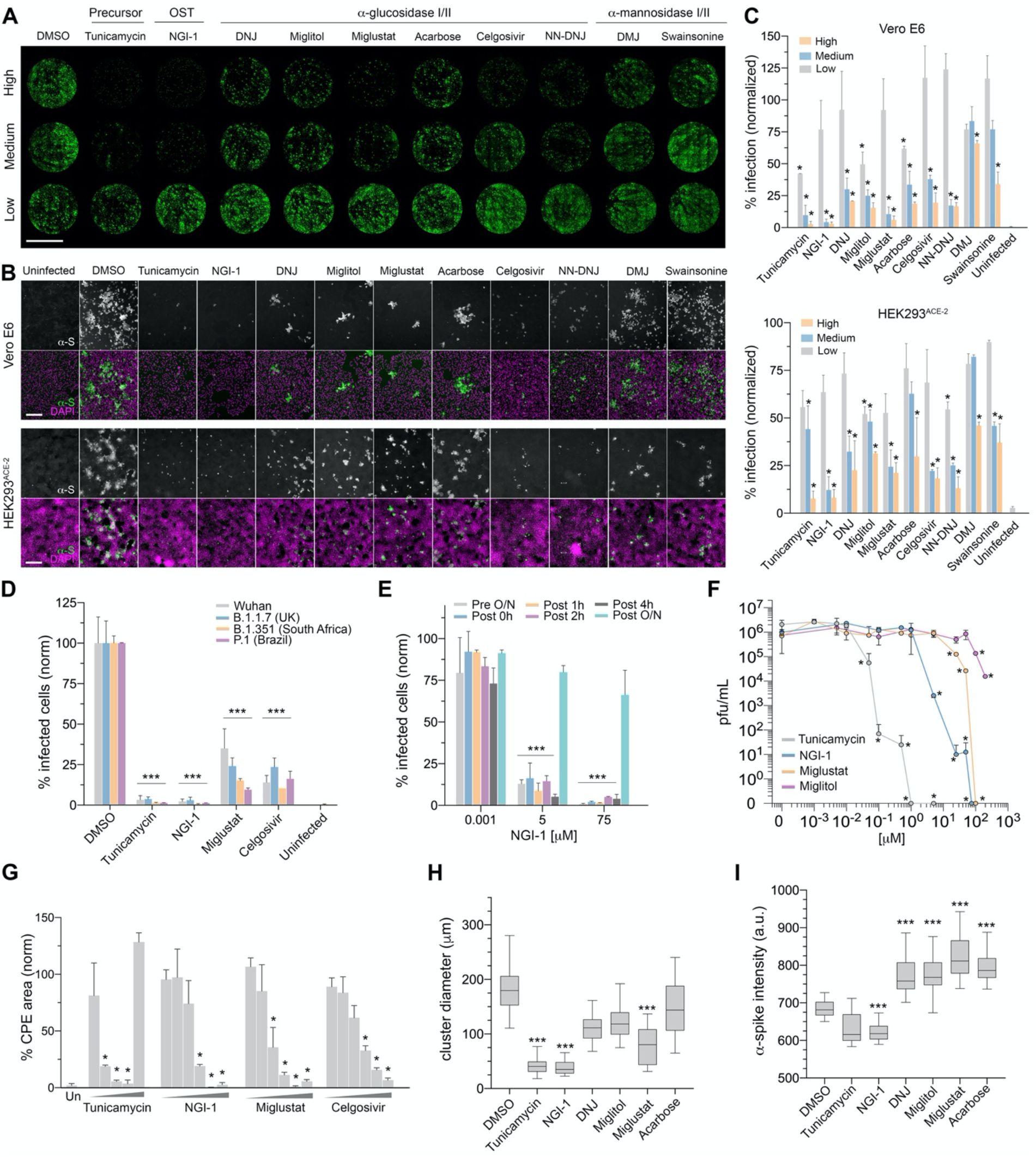
Glycosylation inhibitors reduce the spread of SARS-CoV-2 infection. (**A**) Representative whole-well scans of confluent Vero E6 monolayers treated with either high, medium or low dose (see below) inhibitors and infected with SARS-CoV-2 (MOI=0.05, 24 hours). Cells immunostained using anti-spike antibodies (green). Scale bar 5 mm. (**B**) Representative images of Vero E6 in ‘A’ treated with high dose inhibitors or HEK293^ACE-2^ cells (HEK infected at MOI=0.1, 24 hours); anti-spike (grey), merge (green) with DAPI (magenta). Scale bars 200μm. (**C**) Percentage of infected (anti-spike positive) cells in ‘A-B’; normalized to DMSO controls; two technical replicates; two biological replicates. Inhibitor concentrations low, medium to high (μM): tunicamycin (0.001, 0.01, 0.05); NGI-1 (0.001, 5, 75); DNJ (0.01, 60, 120); Miglitol (0.01, 50, 200); Acarbose (0.01, 100, 300); Miglustat (0.01, 50, 100); Celgosivir, Swainsonine, NN-DNJ and DMJ (0.01, 25, 100). (**D**) Percentage of infected Vero E6 cells pre-treated with high dose inhibitors and infected with either the Wuhan (Liverpool) isolate, variant B.1.1.7, variant B.1.351 or variant P.1; normalized to DMSO controls. (**E**) Percentage of infected Vero E6 cells (MOI=0.05, 24 hours) treated with inhibitors at different time points in relation to inoculation (pre-incubation overnight; 0 hours co-incubation; 1, 2, 4 hours and overnight post-incubation) with 0.001, 5 or 75 μM NGI-1; normalized to DMSO controls. (**F**) Viral titres in Vero E6 supernatants treated with inhibitors and infected with SARS-CoV-2 (MOI=0.001, 48 hours); dots indicate mean values, two technical replicates, two biological replicates. (**G**) Area percentage covered by dead Vero E6 cells treated with inhibitors (low to high dose) and infected with SARS-CoV-2 (MOI=0.001, 48 hours); normalized to DMSO controls; two technical replicates, two biological replicates. (**H**) Box plot of cluster diameter (μm) of infected Vero E6 cells in ‘A-B’ treated with high dose inhibitors; horizontal lines represent mean values, whiskers represent ±CI. (**I**) Fluorescence intensity of anti-spike antibodies in infected Vero E6 cells in ‘A-B’ treated with high dose inhibitors. Bars indicate mean values; error bars represent +S.D.; asterisks indicate significance.

We then hypothesized that virions assembled in cells with altered *N*-glycosylation may present abnormal spike *N*-glycosylation profiles and consequently become less infectious. As a model, we used virions produced in Vero E6 cells treated with 5 μM NGI-1. This treatment was previously observed to greatly reduce the proportion of infected cells while allowing the formation of fewer infective viral particles (**Fig. 2**). Supernatants from infected NGI-1-treated cells contained on average 392-fold less infective viral particles than untreated cells (**Fig. 3A**). However, quantification of total viral RNA (i.e., infective and non-infective particles) showed only 82-fold reduction (**Fig. 3B**). This imbalance between total and infective particles (4.8-fold) may account for virions incapable of infecting host cells as a result of missing or expressing aberrant protein *N*-glycans. To validate the hypothesis, supernatants were used to infect cells normalizing the virus inoculum based on total viral RNA. Virions released by NGI-1-treated cells still produced lower infections (4.4-fold), evidencing the existence of a sub-population of defective virions unable to infect subsequent cells (**Fig. 3C**). Cells pre-treated with NGI-1 showed an even stronger reduction when inoculated with virions produced in NGI-1-treated cells. To confirm that the absence surface *N*-glycans renders viral particles non-infective, we incubated intact SARS-CoV-2 virions with Peptide-N-Glycosidase F (PNGase F) prior to infection (**Fig. 3, D to H**). PNGase F removes all types of *N*-linked oligosaccharides from glycoproteins, except those with core a1,3-fucose. Under this condition, the enzyme is only expected to cleave accessible *N*-glycans from the surface exposed viral glycoproteins spike, membrane (M) and envelope (E). Overall protein deglycosylation was confirmed by lectin blotting (**Fig. S5A**), and by analyzing the mobility of spike and M proteins by western blotting (**Fig. 3D**). Mock-treated (P-) and control (C) virions showed a full-length spike band (S0; ~250 kDa) and four more of higher apparent molecular mass likely corresponding to aggregates resistant to denaturing, multimeric or more glycosylated forms. In virions treated with PNGase F (P+), all bands significantly downshifted due to the loss of *N*-glycans. In addition, two M bands were detected in control samples corresponding to unglycosylated (~25 kDa) and glycosylated (~30 kDa) forms, with the latter collapsing to ~25 kDa upon digestion. Remarkably, deglycosylated virions (P+) were unable to infect host cells, while cells inoculated either with P-or C presented infection levels comparable to those inoculated with non-purified supernatant virions (SN) (**Fig. 3, F to H**). Quantification of total viral RNA (**Fig. S5B**) and infectious particles in cell supernatants (**Fig. S5C**) confirmed that P+ did not develop SARS-CoV-2 infection. Cells inoculated with heat-inactivated PNGase F-treated virions (P_IN_) or co-inoculated with PNGase F (P_T0_) produced normal infections. We then used liquid chromatography coupled to high-resolution tandem mass spectrometry (LC-HR-MS/MS) to identify the spike *N*-glycosylation sites whose glycans were removed by the enzyme. By looking at the conversion of asparagine into aspartate after PNGase F deglycosylation, we found five spike peptides containing seven modified sites (i.e., N234D, N282D, N331D, N343D, N801D, N1158D and N1173D) (**Fig. 3E; Data S2 to S8**). Although other sites could be deglycosylated, removal of the complex-type *N*-glycans on sites N331 and N343 (both within the RBD) is likely to be the cause of P+ infection failure, as suggested in previous studies (*9, 10, 14*). Lastly, deglycosylation of cell surface glycoproteins prior to virus inoculation did not affect the infection outcome (**Fig. 3H and Fig. S5D**), suggesting that host surface *N*-glycans (i.e., susceptible to PNGase F digestion) play no role in initial virus invasion as previously suggested(*28*).

**Fig. 3.**
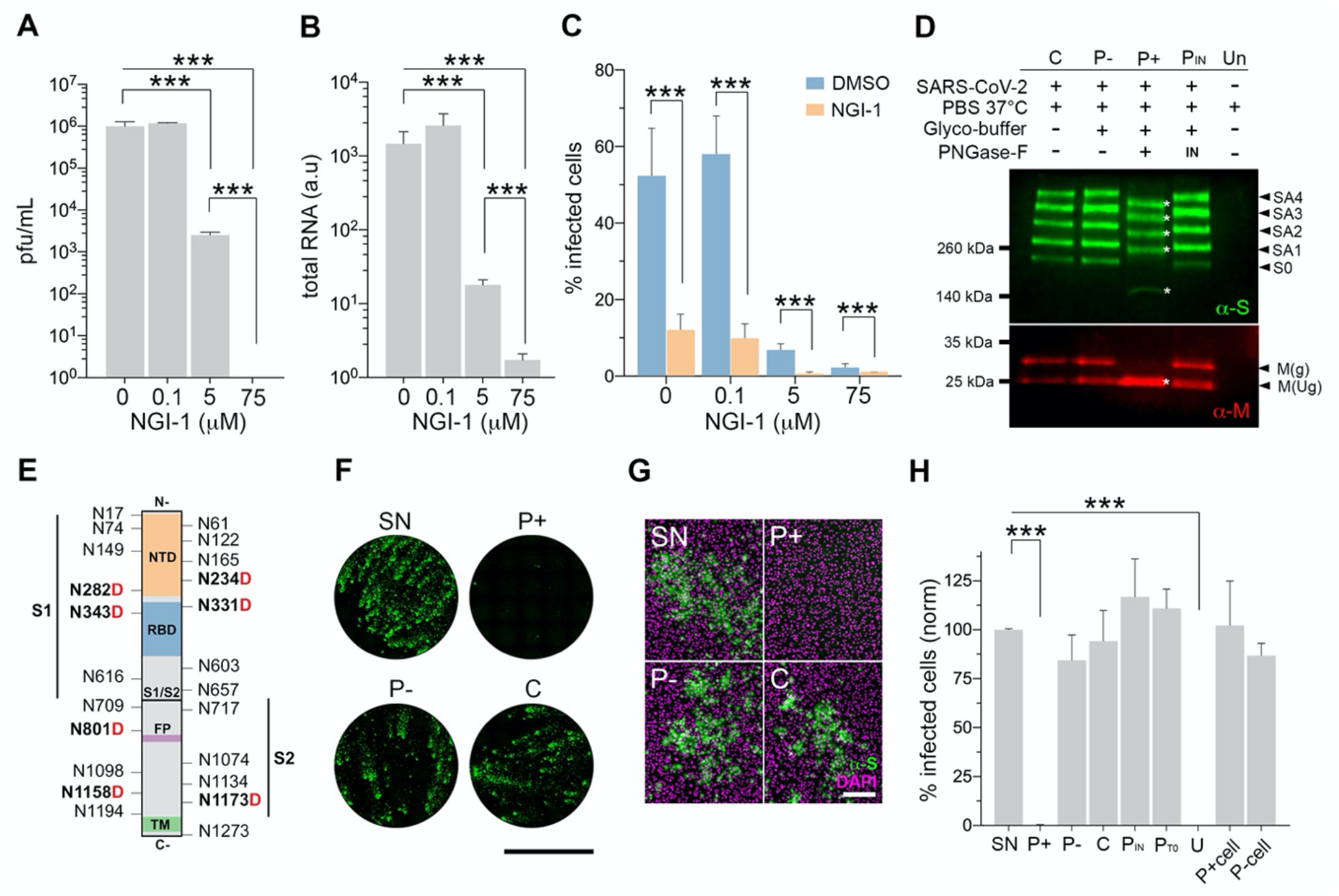
SARS-CoV-2 surface *N*-glycans are essential for infection. (**A**) Infective particles in infected Vero E6 supernatants (MOI=0.001, 48 hours) treated with 0, 0.1, 5, 75 μM NGI-1. (**B**) Total SARS-CoV-2 RNA in supernatants in ‘A’. (**C**) Proportion of infected Vero E6 cells treated with 0, 0.1, 5, 75 μM NGI-1 and infected with virus from either untreated (DMSO) or 5 μM NGI-1-treated cell supernatants (MOI=0.05, 24 hours); inoculum normalized by total viral RNA. (**D**) Western blot using anti-spike (green) and anti-M (red) antibodies on lysed native purified virions (C), mock treated (P-), PNGase F-treated (P+), treated with heat-inactivated PNGase F (P_IN_) or supernatant from uninfected cells (Un). Treatment legend (top), apparent molecular mass (left), band identity (right): full length spike (S0), spike aggregates (SA), glycosylated (g) and unglycosylated (Ug) M protein; asterisks highlight downshifted bands. (**E**) Spike protein schematic highlighting all *N*-glycosylation sites and asparagines (N) converted to aspartic acid (bold red ‘D’) after PNGase F deglycosylation as found by mass spectrometry; spike subunits (S1, S2); N-terminal domain (NTD); receptor binding domain (RBD); fusion peptide (FP); transmembrane domain (TM). (**F**) Whole-well scans of confluent Vero E6 cells infected with SARS-CoV-2 supernatant (SN) or purified virions analyzed in ‘D’ (MOI=0.05, 24 hours); anti-spike antibodies (green), scale bar 5 mm. (**G**) Representative images of Vero E6 cells in ‘F’; anti-spike (green) merged with DAPI counterstain (magenta). Scale bar 200μm. (**H**) Percentage of infected cells in ‘F’ including purified virions co-inoculated with active PNGase F (P_T0_) and cells either treated (P+_cell_) or untreated (P-_cell_) with active PNGase F prior to inoculation with native virions; normalized to SN. Bars indicate mean values, error bars show +S.D., asterisks indicate significance (*p*<0.001).

In summary, we investigated the essentiality of host *N*-glycosylation pathway and SARS-CoV-2 *N*-glycans in relation to infection. Our data suggest that inhibiting host *N*-glycosylation does not prevent virus recognition and invasion in first place. Instead, we propose that the consequent *N*-glycosylation alterations in newly synthesized viral proteins negatively impact the production of new infective virions. Cells with altered *N*-glycosylation release fewer virions with missing or aberrant spike *N*-glycans that are essential for invasion. Overall, altering host *N*-glycosylation leads to a reduction of the spread of SARS-CoV-2 infection, and the earlier the pathway is halted the more effective this reduction becomes (**Fig. 4**). The evidence generated in this study allows further research on developing ways to exploit *N*-glycosylation to treat COVID-19. Some of the inhibitors tested here (i.e., Miglustat, Miglitol, Acarbose and Celgosivir) are drugs in trials or FDA-approved, used to treat unrelated human diseases and could be considered for repurposing if found effective in trials, alone or in combination. Further new treatments may consist in blocking essential spike *N*-glycans using antibodies or lectins to reduce the spread of the infection(*29*). Concerns are currently arising about the constant generation of new SARS-CoV-2 variants that can escape vaccine-derived antibody neutralization. However, our results show evidence that infection by SARS-CoV-2 variants B.1.1.7, B.1.351 and P.1, three of the most common worldwide, is equally affected by glycosylation inhibitors. This brings hope for a standalone therapy and may also maximise the efficacy of current vaccines, as partially glycosylated viruses will be more vulnerable to neutralising anti-spike antibodies(*14*). Importantly, approaches targeting essential *N*-glycans may offer an advantage over protein-based ones as *N*-glycosylation is a more conserved post-translational modification less favourable to mutation. In fact, no mutations concerning *N*-glycosylation sites have been identified to date(*30*). Altogether, the study of *N*-glycosylation in relation to viral diseases may offer new opportunities to fight, not only SARS-CoV-2, but future coronavirus outbreaks.

**Fig. 4.**
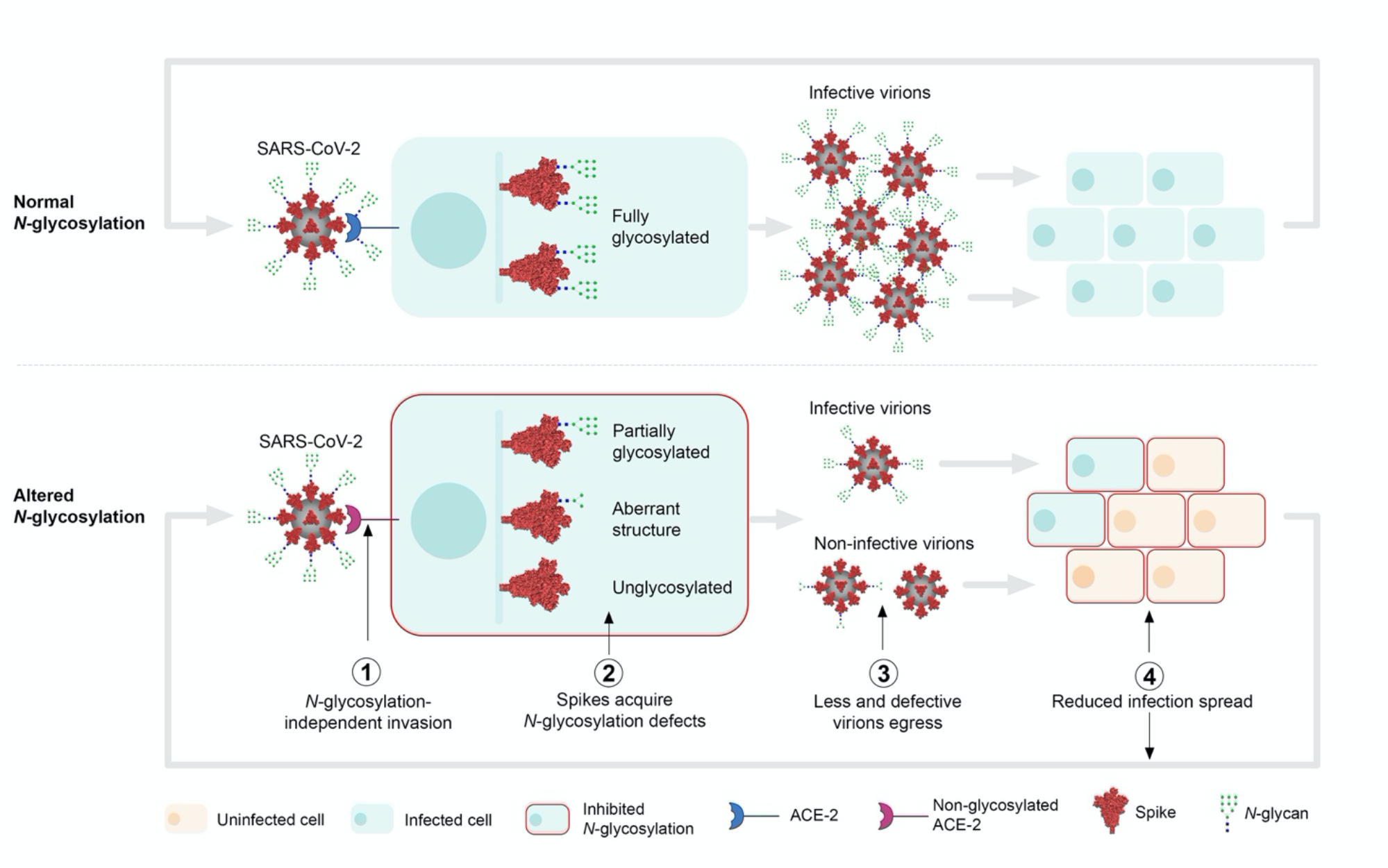
Schematic of the roles of *N*-glycosylation in SARS-CoV-2 infection. Initially, SARS-CoV-2 virions can invade the host cell regardless of its *N*-glycosylation status (i.e., inhibited *N*-glycosylation, deglycosylated ACE-2) (**1**). Once it starts replicating, unlike in untreated cells where spike proteins get fully *N*-glycosylated (66 sites per trimer), cells with inhibited *N*-glycosylation (e.g., drug-treated or glycosylation gene downregulated) will produce spike proteins presenting partial glycosylation (<66 glycosylated sites), aberrant glycan structures and/or no glycosylation at all (**2**). Improperly glycosylated spikes may misfold, accumulate in the ER and/or degrade, leading to egression of less and defective virions (**3**). These virions become less or non-infective to neighbouring cells as they lack spike *N*-glycans essential for invasion (**4**). Replication in these cells with inhibited *N*-glycosylation will generate again less and non-infective virions (**2-4**), therefore amplifying the inhibition of the spread of the infection exponentially. Glycan structures represented using CFG nomenclature; spike and virion glycan structures are representative only; blue vertical bar represents the secretory pathway.

## Acknowledgements

We would like to thank Prof. James D. Bangs, Dr. Spyridon Stavrou, Dr. Thomas Crozier and Dr. Lee R. Haines for critical reading of the manuscript and helpful suggestions.

## Author Contributions

ACS and AAS conceptualized and designed the study. ACS, ARR, EH, CCE, BG, IE and TZ performed the experiments. ACS, ARR, EH, EIP, GLH, ICA, TZ and AAS analyzed, curated and validated data. ACS wrote the paper with input from all the authors.

## Materials and Methods

### Cell and SARS-CoV-2 culture

Vero E6 cells (C1008; green African monkey kidney cells) were obtained from Public Health England and cultured in DMEM medium (Lonza) supplemented with 10% foetal bovine serum (FBS; Gibco) and 50 μg/mL gentamicin (Sigma) at 37°C and 5% CO_2_. HEK293^ACE-2^ cells (lentiviral expression of human angiotensin-converting enzyme 2) were cultured in DMEM supplemented with 10% FBS, 1mM sodium pyruvate (Gibco) and 2 μg/mL puromycin (Invivogen) at 37°C and 5% CO_2_. Cells were regularly passaged (passage 2 to 30) by trypsinization when >80% confluence was reached. SARS-CoV-2 isolate SARS-CoV-2/human/Liverpool/REMRQ0001/20 20, isolated from naso-pharyngeal swab from a patient in March 2020, was passaged four times in Vero E6 cells (*31*). Experimental cells were infected by incubation with SARS-CoV-2 (passage 4 in Vero E6) supernatant containing 2 x 10^7^ pfu/mL in DMEM 2% FBS for 30 minutes at 37°C and 5% CO_2_, at variable MOIs (see below). SARS-CoV-2 variants B.1.1.7, B.1.351 and P.1 were obtained from BEI Resources.

### siRNA knockdowns

A total of 2 x 10^5^ Vero E6 or HEK293^ACE-2^ cells were seeded 24 hours before transfection with one of the following siRNAs pools (Qiagen SMARTpool): siSTT3A (#GS3703), siSTT3B (#GS201595), siGANAB (#GS23193) or siMGAT1 (#GS84061). Double knockdown of STT3A and STT3B was performed by transfecting equal amounts of siSTT3A and siSTT3B pools. Nontargeting control (siNT) was obtained from Dharmacon. 4μL of 100μM siRNA pool were combined with 6μL Lipofectamine RNAiMax (Invitrogen) in 200μL Opti-MEM (Gibco) and incubated for 20 minutes at RT before being added dropwise to the cells. Cells were flushed out with fresh medium 4 hours after transfection. For siGANAB and siMGAT1, 2 hits of siRNA over a 120-hour incubation period were performed while for siSTT3A, siSTT3B and siSTT3A+siSTT3B, 3 hits of siRNA over a 168-hour period were needed. The day before optimum knockdown (96 hours for 2 hits, 144 hours for 3 hits), cells were trypsinized and 2 x 10^4^ cells were seeded for each condition onto 96-well optical plates (Greiner) in quadruplicate and infected the following day. Infections were carried out by incubation with SARS-CoV-2 P4 supernatant at either MOI=0.05 (Vero E6) or MOI=0.1 (HEK293^ACE-2^) for 30 minutes at 37°C and 5% CO_2_, washing and incubation in DMEM 2% FBS 37°C and 5% CO_2_ for 24 hours.

### *N*-glycosylation inhibitors

Cells were treated with either DMSO or several *N*-glycosylation inhibitors (from DMSO stock solutions, DMSO final concentration never exceeding 2%) in DMEM 2% FBS. Inhibitors used: Tunicamycin (Sigma), NGI-1 (Sigma), DNJ (Cambridge Bioscience), Miglitol (Stratech Scientific), Miglustat (Cambridge Bioscience), Acarbose (Fisher Scientific), NN-DNJ (Cambridge Bioscience), Celgosivir (Sigma), DMJ (Tebu-Bio) and Swainsonine (Cambridge Bioscience). Cells were then infected by incubation with SARS-CoV-2 P4 supernatant with the corresponding inhibitor for 30 minutes at 37°C and 5% CO_2_, at either MOI=0.001 (Vero E6 for plaque assay), MOI=0.05 or MOI=0.1 (Vero E6 and HEK293^ACE-2^ for immunofluorescence assay, respectively). Cells were washed and left in DMEM 2% FBS and *N*-glycosylation inhibitor at 37°C and 5% CO_2_ for 48 hours (Vero E6 plaque assay) or 24 hours (immunofluorescence). For toxicity assays, cells in 96-well plates were treated with variable concentrations of inhibitors in DMEM 2% FBS in triplicate and after 48 hours, AlamarBlue 10X solution (Invitrogen) was added and incubated for 4 hours at 37°C before measuring well fluorescence intensity at 590 nm using a FLUOstar Omega plate reader (BMG Labtech).

### Plaque assay

Vero E6 were cultured in DMEM 10% FBS in 24-well plates and allowed to reach confluency overnight. Cells were incubated with 0.1 mL serially diluted supernatant containing SARS-CoV-2 for 30 minutes at 37°C and 5% CO_2_ before being left in 1.5mL overlay (DMEM, 2% FBS, 1.2% cellulose microcrystalline (Sigma)) at 37°C and 5% CO_2_ for 72 hours. Cells were fixed in formaldehyde for 30 minutes, washed and stained with 0.1% crystal violet 20% ethanol solution for manual plaque counting.

### RT-qPCR RNA quantification

Total RNA was extracted from supernatants of SARS-CoV-2-infected Vero E6 cells using the RNeasy kit (Qiagen) following the manufacturer’s protocol. 4μL purified RNA were subsequently used for RT-qPCR reaction using the Luna Universal One-Step RT-qPCR kit (New England Biolabs) and 0.4μM of each primer: CoV2-N-F (CACATTGGCACCCGCAATC) and CoV2-N-R (GAGGAACGAGAAGAGGCTTGAC). Reactions were set and analyzed in an Agilent Mx3005P machine as follows: 55°C for 10 minutes, 95°C for 1 minute, 40 cycles of 95°C for 10 seconds followed by 60°C for 30 seconds; with final melting curve.

### Immunofluorescence assay

Cells cultured in optical plates were fixed in 4% formaldehyde (Pierce) for 30 minutes at RT, washed in PBS and permeabilized in 0.5% saponin for 10 minutes. Cells were then blocked in 1% bovine serum albumin (BSA) PBS for 45 minutes and further incubated in blocking solution containing anti-SARS-CoV-2 spike protein subunit 2 (Abcam) 1:1000 dilution for 1 hour at 37°C. Blocking solution containing DAPI 1μg/mL and Alexa Fluor 488 Plus-conjugated anti-mouse IgG (Thermofisher) was then incubated for 1 hour at RT. Cells were washed and left in PBS for imaging. Imaging was carried out using a Zeiss LSM800 confocal laser scanning microscope by tile-scanning entire plates conserving identical settings between comparable samples. Images were analyzed using Fiji (ImageJ) by thresholding to select for true signal and measuring relative cell area coverage in DAPI channel (total number of cells) and anti-spike area or mean fluorescence intensity (infected cells). Infection rates given as normalized infected area (anti-spike-positive) over total cell area (DAPI).

### PNGase F deglycosylation of intact SARS-CoV-2 virions

2mL SARS-CoV-2 P4 supernatant containing 2 x 10^7^ pfu/mL was purified using an Amicon Ultra column (MWCO 100kDa; Merck) by centrifugation and washing in PBS at 2,000 x *g*. 40 μL purified supernatant were combined with 2,500 units recombinant glycerol-free PNGase F and Glycobuffer 2 (New England Biolabs) following the manufacturer’s protocol for non-denaturing digestion and incubated at 37°C for 5 hours. Parallel controls included incubation of purified virions with or without PNGase F and Glycobuffer, incubation with heat-inactivated (75°C for 10 minutes) PNGase F, and incubation with PNGase F and Glycobuffer during virus inoculation of host cells for 30 minutes. After incubation, samples were either directly used for infection or combined with RIPA buffer supplemented with protease inhibitors to generate lysates for blotting. Alternatively, Vero E6 cells were pre-incubated with or without 100,000 units/mL PNGase F in serum-free OptiMEM for 5 hours at 37°C and 5% CO_2_, before being either inoculated with SARS-CoV-2 or surface-biotinylated. To biotinylate surface proteins, cells were washed in PBS pH=8, incubated with EZ-link NHS-biotin (Thermofisher) 2.5mM for 30 minutes on ice, washed in PBS 100mM glycine and lysed in RIPA buffer and protease inhibitors as above.

### Western, lectin and streptavidin blotting

siRNA cell lysates were obtained at either 120-hours (2 hits) or 168-hours (3 hits) by washing cells in cold PBS and incubation in RIPA buffer with 1% phosphatase and protease inhibitor cocktail (ThermoScientific). Protein concentration was determined using Precision Red (Cytoskeleton) and 60μg of protein were separated using 4-15% SDS-PAGE gels and transferred onto nitrocellulose membrane. Alternatively, virion lysates were run in 4-12% SDS-PAGE and dry-transferred onto PVDF membranes. Membranes were blocked in either 5% milk or 2% BSA in Tris-buffered saline or PBS, 0.1% Tween-20 (TBS-T) for 30 minutes and incubated with their corresponding primary antibodies at 4°C overnight. Primary antibodies used: anti-STT3A (HPA030735, Cambridge Bioscience), anti-STT3B (A15574, Universal Biologicals), anti-GANAB (A13851, Universal Biologicals), anti-MGAT1 (CSB-PA013773ESR1Hu, 2B Scientific), anti-GAPDH (10494-1-AP, ProteinTech), anti-spike S2 (Abcam) and anti-M (. Blots were then incubated with anti-rabbit IgG conjugated with horseradish peroxidase (Cell Signalling) or Alexa Fluor Plus 800 (Thermofisher), or anti-mouse IgG conjugated with Alexa Fluor Plus 680 (ThermoFisher) for 1 hour. Blots were visualized on a BioRad ChemiDoc™ Touch Gel Imaging System using enhanced chemiluminescence substrate (for horse radish peroxidase secondary) or on a LI-COR Odyssey Fc (for fluorescence). Virion lysate blots were then incubated with 5 μg/mL concanavalin A FITC-conjugated in 1% BSA 0.05% Tween-20 PBS for 30 minutes. GAPDH or nigrosine staining were utilized as loading control. Streptavidin blots were incubated in streptavidin-Alexa Fluor 555 (Thermofisher) 5μg/mL in PBS 0.1% Tween20, 0.02% SDS for 30 minutes at RT. Lectin and streptavidin blots were imaged using a Thermofisher iBright FL1500 system.

### *N*-glycoprofiling

Either siRNA- or inhibitor-treated cells were harvested by trypsinization, washed in cold PBS, manually counted and snap frozen. A total of 2×10^6^ (Vero E6) or 3×10^6^ (HEK293^ACE-2^) cells were used for the analyses, alongside Ludger IgG as positive and water as negative controls for insolution *N*-glycan release, labelling and clean-up. Samples were denatured at 100°C with SDS and dithiothreitol before incubation overnight with PNGase F in NP-40 buffered solution. The released glycans were then converted to aldoses with 0.1% formic acid, filtered through a protein binding membrane and dried. *N*-glycans were then labelled with procainamide, cleaned up in water, dried and reconstituted in water. Samples were analysed by HILIC-UPLC using an ACQUITY UPLC BEH-Glycan 1.7 mm, 2.1×150mm column at 40°C on a Thermo Scientific UltiMate 3000 UPLC with a fluorescence detector (excitation = 310 nm; emission = 370nm) controlled by HyStar software v3.2. Gradient conditions were: 0 to 53.5 min, 24 to 49 % Buffer (A) at a flow rate of 0.4 mL/min; 53.5 to 55.5 min, 49 to 100% A, 0.4 to 0.2 mL/ml; 55.5 to 57.5 min, 100% A at 0.2 mL/min; 57.5 to 59.5 min, 100 to 24% A at 0.2 mL/min; 59.5 to 65.5 min, 24% A at 0.2 mL/min; 65.5 to 66.5 min, 24% A from 0.2 to 0.4 mL/min; 66.5 to 70 min 24% A at 0.4 mL/min. Buffer A was 50 mM ammonium formate; Buffer B was acetonitrile (Acetonitrile 190 far UV/gradient quality). Samples were injected in 25% aqueous/75% acetonitrile; injection volume 25 μL. Ludger procainamide labelled glucose homopolymer, A2 & A3 mix, FA2 mix and Man mix were used as standard and as external calibration for GU allocation. All standards were run in the same injection volume and conditions as the samples. Thermo Scientific Chromeleon software v7.2.1 was used to allocate GU values to peaks. When software did not split peaks that were not fully resolved, areas were merged together across peaks. Glycan analyses were performed by Ludger Ltd (*32-34*).

### Trypsin digestion for proteomic analysis

Purified virions were subjected to PNGase F treatment (P+; sample#3) or heat-inactivated PNGase F treatment (P_IN_; sample#4) as described above. The positive controls (for viral proteins) included purified virions (C; sample#1) and PNGase F mock-treatment of purified virions (P-; sample#2). The negative controls (devoid of viral proteins) included DMEM 2% FBS (sample#5) and Vero E6 cell culture conditioned supernatant or secretome in DMEM 2% FBS (sample#6), both purified by size exclusion as described above. Samples were incubated in 80% ice-cold acetone for 1 hour at −20°C for protein precipitation, spun down at 14,000x*g* for 15 min and pellets were air-dried. Peptides were obtained from the via FASP Protein Digestion Kit (#ab270519, Abcam, Cambridge, MA), using trypsin (Proteomics Grade, #T6567, Sigma-Aldrich, St. Louis, MO). Lyophilized intact protein samples #1-6 (141-478 μg protein) were redissolved in 200 μl 100 mM Tris-HCl pH 7.0, containing 8M urea (urea sample solution, USS). Protein digestion was performed with 100 μl of each sample and transferred to a 1.5mL microtube containing 200 μL USS. 100 mM DL-dithiothreitol (catalog #D0632, Sigma-Aldrich) was added to reduce protein disulfide bonds and achieve complete protein unfolding. Samples were vortexed for 30 s and then kept swaying on a nutating mixer (Nutator Mixer Model 117, TCS Scientific Corp.) for 45 min at RT prior to proteome extract digestion protocol. Samples were transferred to a 30-kDa Spin Filter (FASP Kit), previously assembled on a collection tube and centrifuged at 14,000 x *g* for 15 min at RT. 200 μl USS was added to each Spin Filter and further centrifuged for 15 min. Flow-through from the collection tube was discarded and 100 μl of fresh prepared 10 mM iodoacetamide, resuspended in USS, was added to the Spin Filter and samples were subsequently subjected to incubation without mixing for 20 min in the dark at RT for alkylation. After incubation, the Spin Filters were centrifuged at 14,000 x *g* for 10 min at RT. 100 μl of USS was added to the Spin Filter and samples were centrifuged at 14,000 x *g* for 15 minutes. This step was repeated twice. Flow-through was again discarded. 100 μL 50 mM ammonium bicarbonate solution was added to the Spin Filter and sample was centrifuged at 14,000 x *g* for 15 min, at RT. This step was repeated twice. After USS removal by ammonium bicarbonate, Spin Filter containing protein sample was transferred to a new collection tube and 100 μL trypsin was added to the Spin Filter, at an enzyme-to-protein ratio of 1:50. Evaporation was prevented by wrapping the top of the tubes with parafilm. The Spin Filter was incubated at 37°C for 16 h without mixing. 200 μL LC-MS grade 0.1% formic acid was added to stop the proteolysis, and each sample was then centrifuged at 14,000 x *g* for 10 min to obtain the filtrate containing the tryptic peptides. The collection tubes containing the peptides were dried down in a Speedvac concentrator (Savant, Thermo Fisher Scientific) and the volume was adjusted with LC-MS grade 0.1% formic acid (Thermo Fisher Scientific) to a final concentration of 1 μg/μL of tryptic peptides prior to liquid chromatography-high resolution-tandem mass spectrometry (LC-HR-MS/MS) analysis.

### Proteomic analysis by LC-HR-MS/MS

The equivalent of 1 μg microgram of protein digest from each sample (samples #1-6), as measured by NanoDrop One/One Microvolume UV-Vis Spectrophotometer (Thermo Fisher Scientific) was loaded onto a 50-cm μPAC capLC column (PharmaFluidics, Ghent, Belgium) separated by a Dionex Ultimate 3000 RSLCnano (Thermo Fisher Scientific) with 99% solvent A (100% water, 0.1% formic acid) and 1% solvent B (90% acetonitrile, 0.1% formic acid), at a flow rate of 750 nL/min for 5 min. The flow rate was decreased to 300 nL/min at 5.1 min, to start the elution gradient. Solvent B was increased to 22% over 64.9 min, then to 45% solvent B over 15 min. The gradient was increased to 95% solvent B over 5 min and maintained at a plateau for an additional 9 min. The gradient sharply decreased to 1% solvent B over 1 min for equilibration with a flow rate at 750 nL/min, maintained for 10 min, thus ending the 120-min sample runtime. Eluted peptides were ionized with a μPAC Flex iON connect (PharmaFluidics, Ghent, Belgium) attached to a Nanospray Flex Ion Source (Thermo Fisher Scientific) equipped with a nanoESI emitter (FOSSILIONTECH, Madrid, Spain). Data was acquired using a Q Exactive Plus Hybrid Quadrupole-Orbitrap Mass Spectrometer (QE Plus, Thermo Fisher Scientific). The QE Plus analysis parameters were as follows: full MS resolution at 70,000; AGC target of 1e^6^; and scan from 350 to 1400 *m/z* range. Top 10 data-dependent MS^2^ parameters were set to 17,500 resolution, ACG target of 1e^5^, (N)CE: 27, charged exclusion at unassigned, +1, +6-8, +>8 with mass exclusion set to ON. Samples #5 and #6 were run previously to generate an exclusion list. The top 200 ions were collected every 5 min (in a total run of 120 min) to generate a 4,800-ion exclusion list prior to sample injection containing virions (samples#1-4). All samples were run in technical duplicate.

### Bioinformatic analysis of proteomic data

Proteomic data analysis was initially performed using Proteome Discoverer (PD) v2.5.0.400 (Thermo Fisher Scientific), with an estimated false discovery rate (FDR) of 1%. Common contaminants such as trypsin autolysis fragments, human keratins, and protein lab standards, were included as well as in-house contaminants, which may be found in the cRAP contaminant database (*35*). SARS-CoV-2 database (36,795 entries) was downloaded in FASTA format on May 18, 2021, from UniProtKB (http://www.uniprot.org/). Additionally, a second SARS-CoV-2 database was added to include the modification of asparagine (N) to aspartic acid (D), resulting from PNGase F treatment (*36*), to illustrate the conserved *N*-glycosylation sequon N-X-S/T, where X is any amino acid except proline. The following parameters were used in the PD analysis: HCD MS/MS; fully tryptic peptides only; up to 2 missed cleavages; parent-ion mass tolerance of 10 ppm (monoisotopic); and fragment mass tolerance of 0.6 Da (in Sequest) and 0.02 Da (in PD 2.1.1.21) (monoisotopic). A filter of two-high confidence peptides per protein were applied for identifications. PD dataset was further processed through Scaffold Q+ 4.8.2 (Proteome Software, Portland, OR) to obtain the protein quantification and further analysis of the MS/MS spectra. A protein threshold of 95%, peptide threshold of 90%, and a minimum of 1 peptide were used for protein validation by manual inspection of each MS/MS spectrum. Proteomics raw data uploaded into PRIDE-EMBL-EBI, internal ID#507100.

## Supplementary Text

### Glycan analyses confirm alterations in protein *N*-glycosylation after genetic and chemical inhibition

To corroborate changes in protein glycosylation upon siRNA or drug treatment, we compared the profiles of procainamide-labeled *N*-glycans from treated and untreated cells using HILIC-UPLC (**Fig S3 and S4; Data S1**). Both cell types appear to express similar glycan profiles, although some species differ in abundance. For example, while the major species in untreated HEK cells were Man5GlcNAc_2_ (M5, peak 19), M6 (peak 30), M7/FA2G1S1 (peak 41), M8 (peak 53), and M9/A3G3S3 (peak 62), Vero cells appear to produce mainly oligomannose glycans and less of the hybrid- and complex-type ones (**Fig S4A**). When STT3-A was knocked out in Vero cells only a slight reduction (26.7%) in the overall levels of *N*-glycans was observed (**Fig S3B**), potentially explaining the absence of protective effect against SARS-CoV-2 (**Fig 1.B-D**). In contrast, both siSTT3-B and siSTT3A+B presented a stronger glycan downregulation (77.4% and 84.7% respectively). Except for a few small structures whose abundances increased (e.g., peaks 6 and 12) in siSTT3-A cells, all siSTT3 cells presented a significant reduction in the abundance of the most common glycans: peaks 2, 4, 19 (M5), 30 (M6), 42 (M7), 53 (M8) and 62 (M9/A3G3S3). The knockdown of GANAB and MGAT1 barely reduced the total amount of glycans (22% and 18%, respectively). siGANAB cells had reduced the major glycans M6, peaks 2 and 14, and completely lost peak 4. siMAGT1 cells accumulated, among others, M5, peaks 4, 14, 23 and 33 while reducing M6, M7, and peak 26. On the other hand, HEK siRNA cells presented *N*-glycan profile modifications slightly different to those seen in Vero cells (**Fig S4B**). siSTT3-A was the only STT3-targeting treatment that significantly reduced glycan levels globally (43%), since siSTT3A+B barely produced any effect (7% reduction). siGANAB and siMGAT1 led to increases of 7.5% and 45.1%, respectively, the latter mainly due to a more pronounced accumulation of M5 compared to that in Vero cells. When either Vero or HEK cells were incubated with glycosylation inhibitors, abnormal *N*-glycosylation patterns were observed compared with that from untreated cells (**Fig S3 and S4**). As expected, incubation of HEK cells with either tunicamycin or NGI-1 significantly reduced the overall amount of *N*-glycans (56.3% and 53.8%, respectively). On the other hand, each α-glucosidase inhibitor modified the *N*-glycan profile in a unique manner; i.e. DNJ reduced the overall glycan levels down to 67.8%, while NN-DNJ, Miglustat, Miglitol, and Celgosivir increased it by 1.0%, 29.5%, 59.1% and 182.6%, respectively. While Miglitol barely modified the relative glycan composition –except for an increase in major peaks 51, 53 and 62, corresponding to the assigned structures FA2G2S1, M8 and M9/A3G3S3, respectively– Miglustat mainly promoted an increase in peaks 2, 13, 46 (A2G2S1), 51 (FA2G2S1), 53 (M8) and 62 (M9/A3G3S3). In addition, NN-DNJ augmented the abundance of a broad range of peaks, including 9, 37 and 46 (A2G2S1) while decreasing the major peaks 13, 53 (M8) and 62 (M9). Celgosivir led to the accumulation of peak 46 (A2G2S1), perhaps at the expense of reducing the amount of M9/A3G3S3 (peak 62). Lastly, we analyzed the *N*-glycosylation profiles of SARS-CoV-2-infected Vero cells either untreated or treated with several inhibitors to determine whether viral infection has a role in modifying protein *N*-glycosylation (**Fig S4**). Infection of untreated cells led to a slight reduction (38.3%) of total *N*-glycans. It did modify the overall glycosylation profile of cells by reducing peaks 2, 6, 9 and 14, while also decreasing the amounts of M6, M8 and M9 structures. Infected cells treated with inhibitors presented even greater glycosylation reductions and profile alterations likely due to the combination of both effects.

### Proteomics to identify deglycosylated spike sites

Using high-resolution mass spectrometry, we identified the *N*-glycosylation sites that were susceptible to PNGase F treatment on the spike protein. To that end, purified virions produced by Vero E6 cells were subjected to treatment with PNGase F alone, which after cleavage converts the Asn (N) residues into Asp (D) (*36*). As negative controls for the N-to-D conversion, we used purified virions treated with heat-inactivated PNGase F, as well as purified virions alone or subjected to PNGase F mock treatment. Following treatment, samples were subjected to trypsin digestion and the resulting peptides were analyzed by LC-HR-MS/MS using a high-resolution 50-cm μPAC capLC C18 column directly coupled to the QE Plus MS. The identified proteins and peptides from SARS-CoV-2 (samples #1-4) following PD and Scaffold Q+ analyses are shown in **Data S2 to S7**. Clusters of spike (S), membrane (M), and nucleocapsid (N) proteins were identified (**Data S2**). Based on the normalized total spectra of each sample, N proteins were the major proteins found in all virion-containing samples, followed by spike and M proteins were of low abundance (**Data S3**). To locate the D residues resulting from PNGase F removal of *N*-glycans from asparagine residues, we used an in-house script that converted the N-to-D in all SARS CoV-2 proteins deposited in the UniProtKB database, including the 22 potential *N*-glycosylation sites in the spike protein. **Data S4** shows all the identified spike proteins in the four samples. We included the complete sequence of the spike proteins highlighting the peptides identified by LC-HR-MS/MS to facilitate the visualization of the location of these peptides in each protein sequence and the protein sequence coverage. While no deglycosylated peptides were detected in positive control samples (purified virions alone and mock-treated purified virions) (**Data S5**), in purified virions treated with PNGase F we detected identified five peptides with N-to-D conversion (DLPQGFSALEPLVDLPIGI**<u>D</u>IT**R, YNE**<u>D</u>GT**ITDAVDcALDPLSETK, FP**<u>D</u>IT**NLcPFGEVF**<u>D</u>AT**R, TPPIKDFGGF**<u>D</u>FS**QILPDPSKPSK, and **<u>D</u>HT**SPDVDLGDISGI**<u>D</u>AS**VVNIQK) (**Data S5**). The MS/MS fragmentation and fragment-ion assignments of these peptides are shown in **Data S8**. Of particular interest, peptide FP**<u>D</u>IT**NLcPFGEVF**<u>D</u>AT**R (residues 329-346) is located within the spike RBD, which is essential for the virus binding to the ACE-2 human receptor. A BLASTp analysis of this peptide sequence showed that in all 5,000 spike protein sequences analyzed (maximum number of sequences allowed by BLASTp search), the sequence is fully conserved, strongly indicating that these two *N*-glycans could be essential for interaction with ACE2. Moreover, we found six peptide sequences in the purified virion sample treated with PNGase F (sample # 3) in which the N residue had not been substituted by D, thus indicating some S protein *N*-glycosylation sites remain unglycosylated (**Data S4**). It is important to underscore that PNGase F does not cleave complex *N*-glycans with a fucose residue α1-3-linked to the reducing GlcNAc linked to the N residue (36, 37). This indicates that most likely the two cleaved complex-type *N*-glycans on N331 and N343 are modified with an α1-6-linked fucose residue. Finally, since part of the M protein appears to be *N*-glycosylated (**Fig. 3D**) we also searched for any potential N-to-D conversion after PNGase F treatment. Intriguingly, we found that all tryptic peptide sequences identified in M were located near the C-terminus of the protein, but we failed to find peptide sequences indicative of an N-to-D conversion (**Data S7**). Of note, the only potential *N*-glycosylation sequon found in 763 M protein sequences analyzed through BLASTp and multiple sequence alignment was located at the N-terminus within an apparent ER signal peptide (SP) (e.g., MSDS**<u>N</u>GT**ITV…, MADS**<u>N</u>GT**ITV…), which could not be confirmed by SignalP 4.1 analysis (http://www.cbs.dtu.dk/services/SignalP-4.1/). Therefore, the shift in migration of M following *N*-glycan removal by PNGase F (**Fig. 3D**) can only be explained if the non-canonical *N*-terminus SP of M proteins is not cleaved after protein synthesis in the ER and the *N*-glycan is eventually attached to it. This suggests that the N-terminus tryptic peptide containing the N-glycan is either further processed (e.g. cleaved or modified with another post-translational modification) or was somewhat altered during sample preparation.

**Fig. S1.**
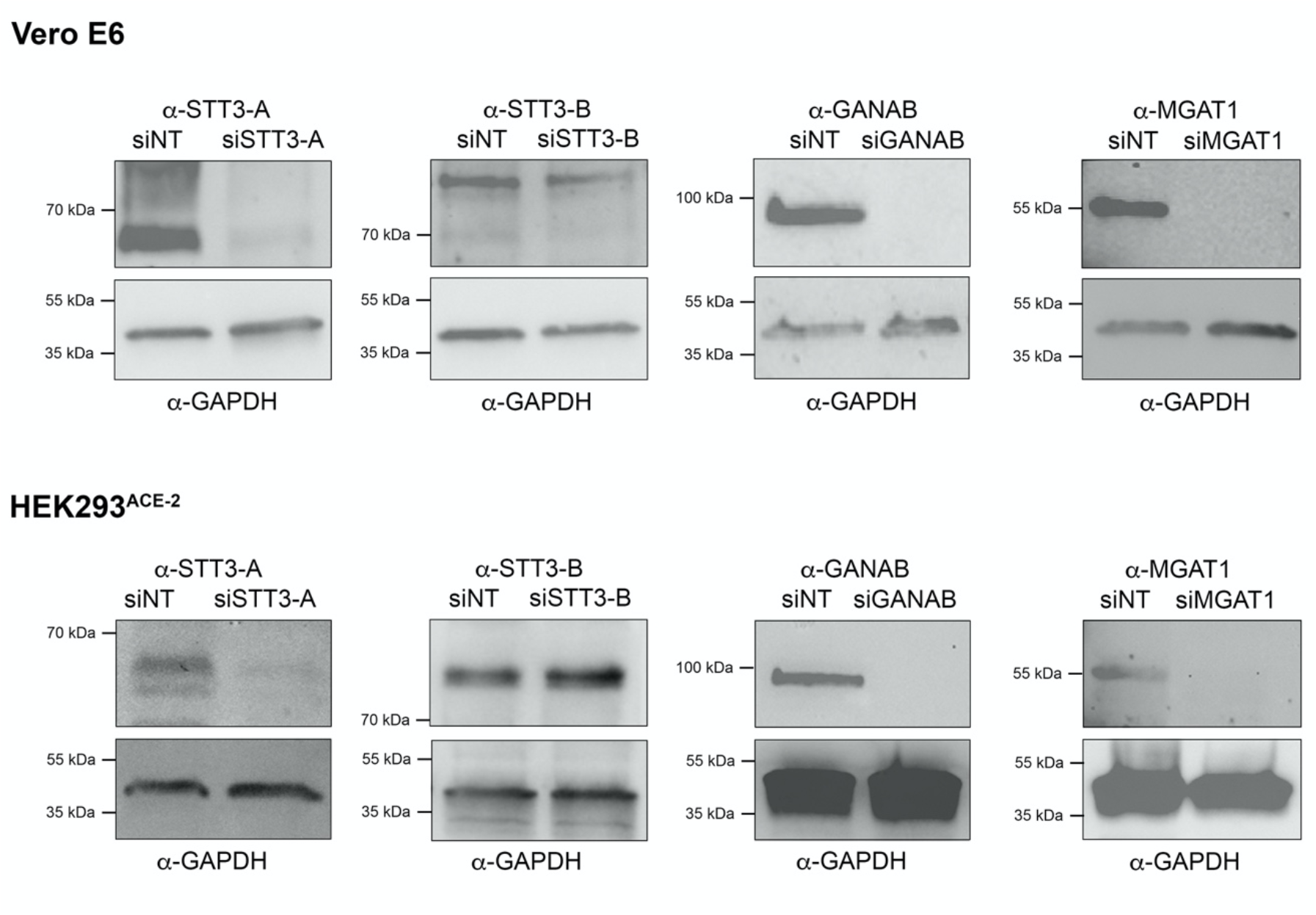
siRNA knockdown validation. Western blotting of lysates from either Vero E6 or HEK293^ACE-2^ cells treated with siRNA targeting glycosylation genes (i.e., STT3-A, STT3-B, GANAB and MGAT1). Lysates probed with specific antibodies against the targeted protein; non-targeted control (siNT). Estimated apparent molecular weights annotated. GAPDH was used as loading control.

**Fig. S2.**
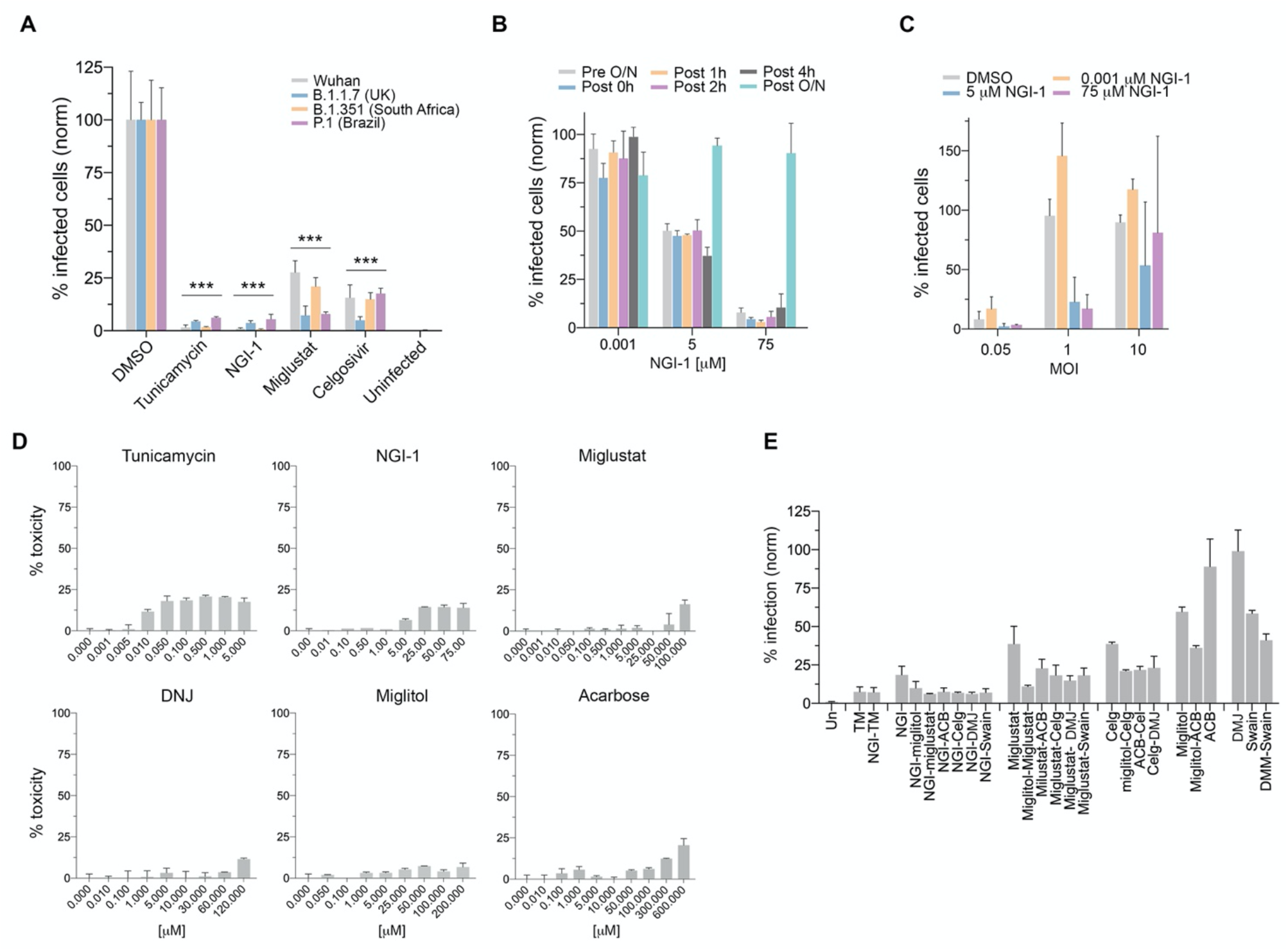
(**A**) Percentage of infected Vero E6 cells pre-treated with high doses of glycosylation inhibitors and inoculated with either the SARS-CoV-2 Wuhan (Liverpool) isolate or the variants B.1.1.7, B.1.351 and P.1; normalized to DMSO controls. (**B**) Percentage of infected Vero E6 cells inoculated with SARS-CoV-2 (MOI=1; 24 hours) and treated with NGI-1 0.001, 5 or 75 μM; normalized to DMSO controls. NGI-1 was added either 16 hours before virus inoculation (Pre O/N), during inoculation (Post 0h), or 1, 2, 4 and 16 hours after inoculation (Post 1h, 2h, 4h, O/N). (**C**) Percentage of infected Vero E6 cells at 24 hours post-inoculation, pre-treated with 0 (DMSO), 0.001, 5 or 75 μM NGI-1 and inoculated with SARS-CoV-2 at MOI= 0.05, 1 or 10. (**D**) AlamarBlue toxicity assays on Vero E6 cells incubated with a range of inhibitors at different doses (μM) for 48 hours. (**E**) Percentage of infected cells pre-treated overnight with combinations of glycosylation inhibitors and inoculated with SARS-CoV-2 (MOI=0.05, 24 hours); normalized to DMSO controls. Bars represent mean values; error bars indicate +S.D.; asterisks indicate significance (*p*<0.001), one-sided *t*-test.

**Fig. S3.**
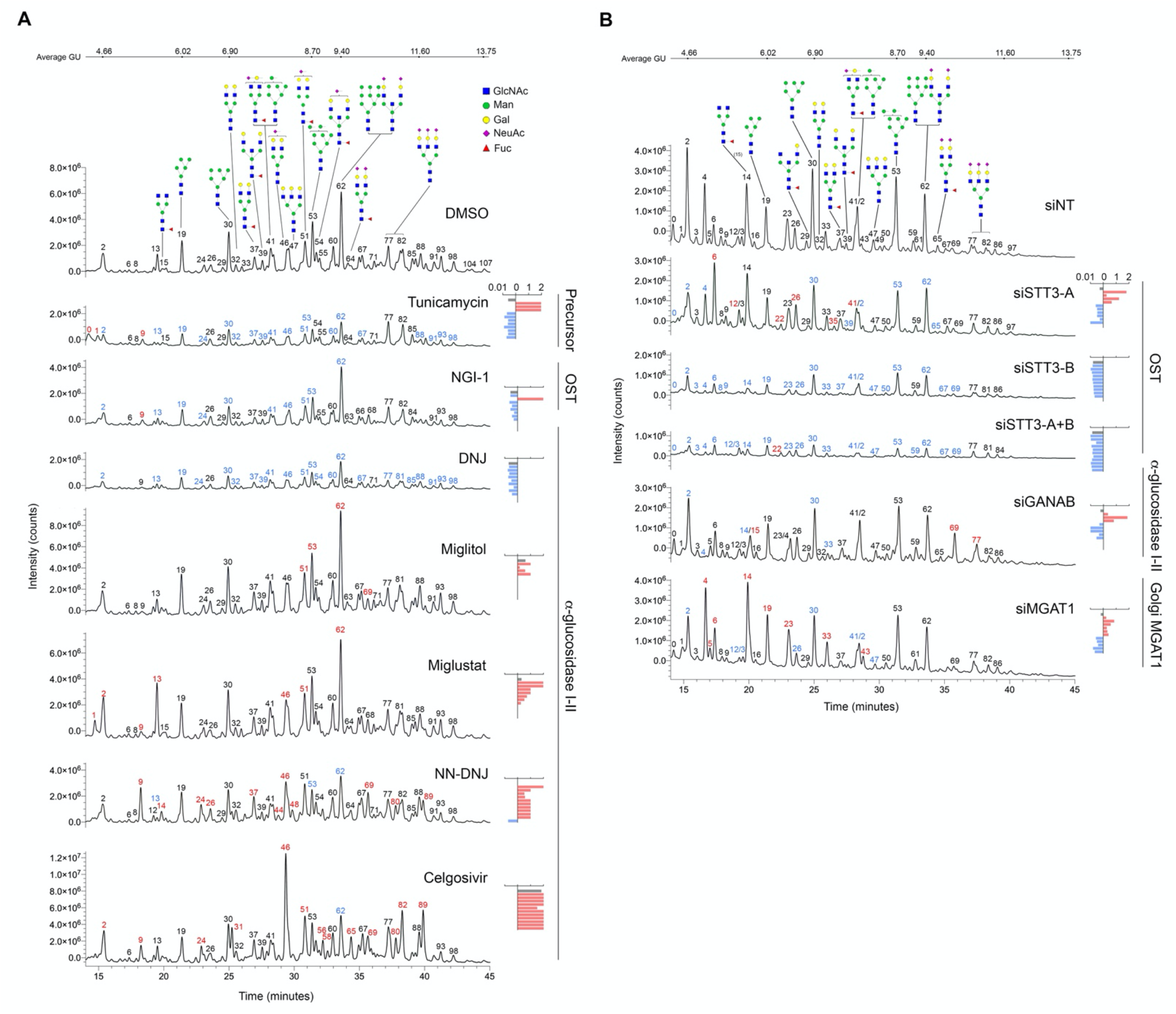
HILIC-UPLC chromatograms showing peak fluorescence intensity (total counts) along time (minutes) of uninfected HEK cells treated with glycosylation inhibitors (**A**), and uninfected Vero cells transfected with siRNAs targeting glycosylation genes (**B**). Control sample (i.e., DMSO, siNT) peaks annotated with possible structures (CFG nomenclature; legend in A) based on glucose units (GU; noted in the top bar). Peaks identified with numbers; colours represent intensity decrease (blue) or increase (red) compared to the control. Bar graphs represent intensity fold change of the sum of all peaks (gray) and individual peaks with the greatest decrease (blue) or increase (red) in intensity compared to the control. The procainamide molecule linked to each glycan is omitted in this figure.

**Fig S4.**
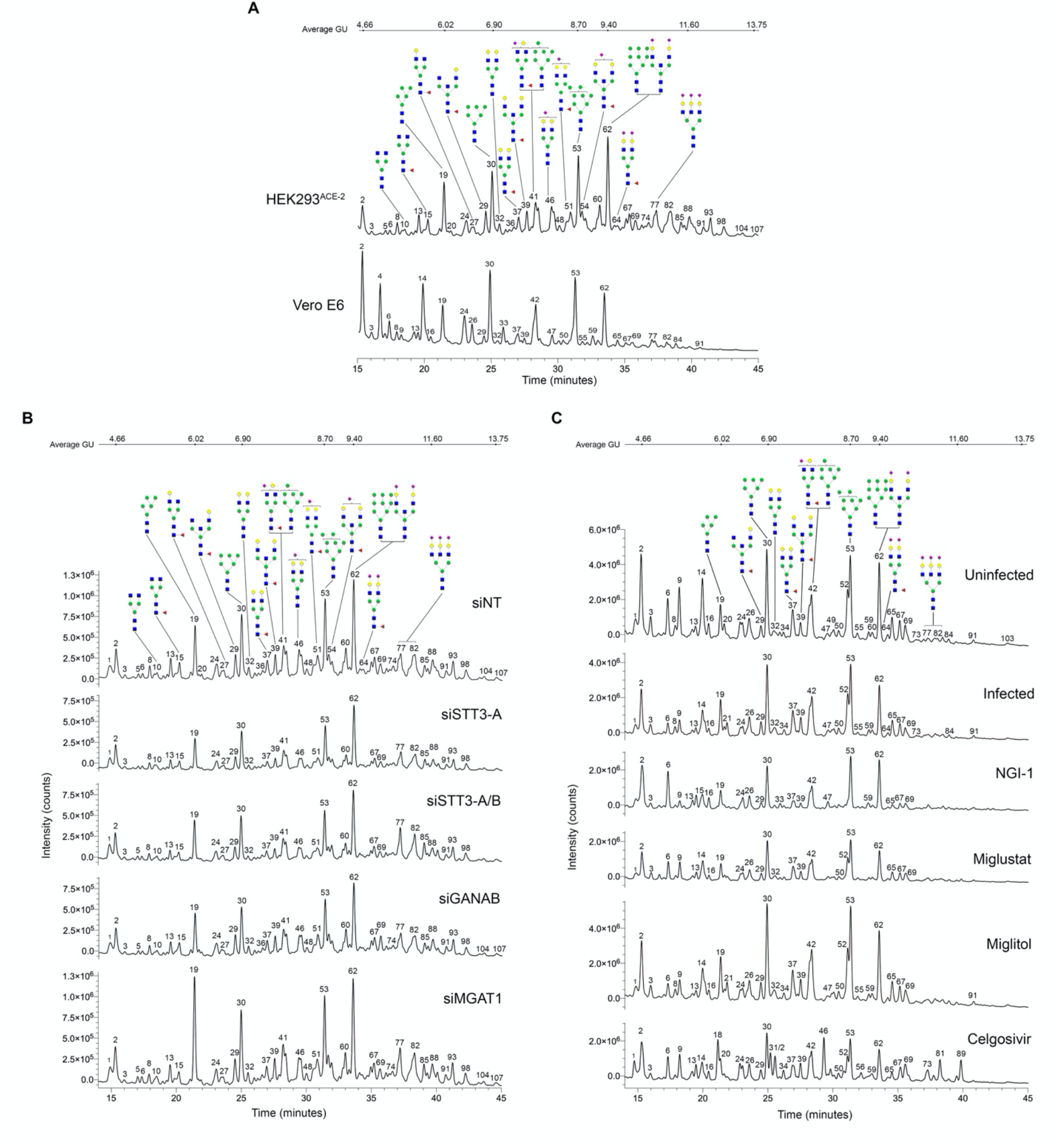
HILIC-UPLC chromatograms plotting fluorescence intensity total counts over time (minutes) of (**A**) untreated HEK293^ACE-2^ and Vero E6 cells; (**B**) HEK293^ACE-2^ cells treated with siRNAs targeting glycosylation genes (STT3-A, STT3A+B, GANAB and MGAT1); (**C**) Vero E6 cells pretreated overnight with glycosylation inhibitors and infected with SARS-CoV-2. Average glucose units (GU) indicated in the top bar; peaks annotated with glycan possible structures annotated on control samples (CFG nomenclature; legend in ‘A’).

**Fig S5.**
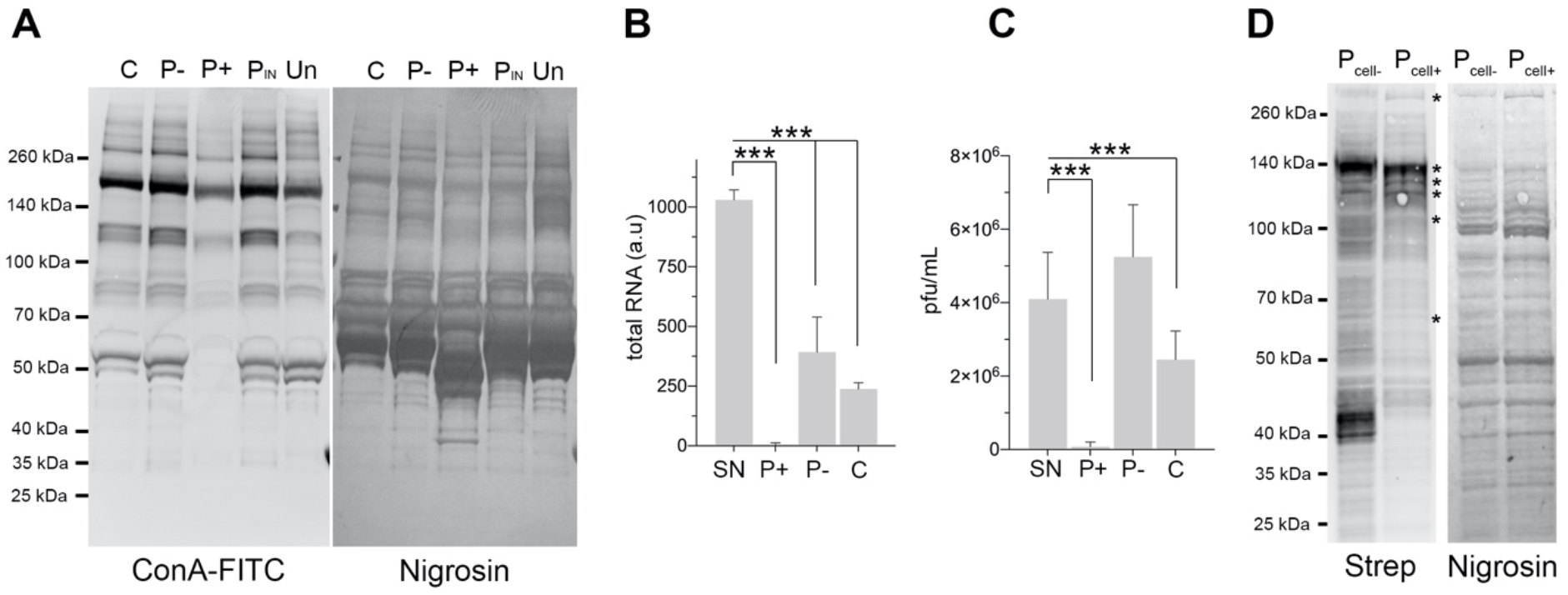
(**A**) Lectin blotting of virion lysates from ‘Fig. 3D’ probed with concanavalin A-FITC (ConA); membrane stained with nigrosin as loading control; apparent molecular weight (left). Although most of the bands recognized by the lectin are likely to be residual serum glycoproteins from the culture media and Vero E6 secretome, there is an overall decrease in recognition after PNGase F treatment. (**B**) Total viral RNA quantification by qRT-PCR in supernatants from ‘Fig. 3E’. (**C**) Virus titres in supernatants from ‘Fig. 3E’ quantified by plaque assay. Bars indicate mean values; error bars show +S.D.; asterisks indicate significance (*p*<0.001), one-sided *t*-test. (**D**) Blotting probed with streptavidin conjugated to Alexa Fluor 555 on lysates of Vero E6 monolayers either untreated (Pcell-) or treated (Pcell+) with PNGase F prior to infection, and surface-biotinylated. Apparent molecular weight (left); nigrosine staining as loading control (right); asterisks indicate biotinylated surface proteins that have downshifted after PNGase F deglycosylation.

## References

1. R. Lu et al., Genomic characterisation and epidemiology of 2019 novel coronavirus: implications for virus origins and receptor binding. Lancet. 395, 565–574 (2020).

2. M. Hoffmann et al., SARS-CoV-2 Cell Entry Depends on ACE2 and TMPRSS2 and Is Blocked by a Clinically Proven Protease Inhibitor. Cell. 181, 271–280.e8 (2020).

3. A. C. Walls et al., Structure, Function, and Antigenicity of the SARS-CoV-2 Spike Glycoprotein. Cell. 181, 281–292.e6 (2020).

4. A. Shajahan et al., Comprehensive characterization of N- and O-glycosylation of SARS-CoV-2 human receptor angiotensin converting enzyme 2. Glycobiology (2020), doi:10.1093/glycob/cwaa101.

5. L. Casalino et al., Beyond Shielding: The Roles of Glycans in the SARS-CoV-2 Spike Protein. ACS Cent Sci. 6, 1722–1734 (2020).

6. A. Bernardi et al., Development and simulation of fully glycosylated molecular models of ACE2-Fc fusion proteins and their interaction with the SARS-CoV-2 spike protein binding domain. PLoS ONE. 15, e0237295 (2020).

7. Y. Watanabe et al., Native-like SARS-CoV-2 Spike Glycoprotein Expressed by ChAdOx1 nCoV-19/AZD1222 Vaccine. ACS Cent Sci. 7, 594–602 (2021).

8. J. Brun et al., Analysis of SARS-CoV-2 spike glycosylation reveals shedding of a vaccine candidate. bioRxiv, 2020.11.16.384594 (2020).

9. Y. K. Choi et al., Structure, Dynamics, Receptor Binding, and Antibody Binding of the Fully Glycosylated Full-Length SARS-CoV-2 Spike Protein in a Viral Membrane. J Chem Theory Comput (2021), doi:10.1021/acs.jctc.0c01144.

10. T. Sztain et al., A glycan gate controls opening of the SARS-CoV-2 spike protein. bioRxiv, 2021.02.15.431212 (2021).

11. R. Henderson et al., Controlling the SARS-CoV-2 spike glycoprotein conformation. Nat Struct Mol Biol. 27, 925–933 (2020).

12. K. M. Bouwman et al., Multimerization- and glycosylation-dependent receptor binding of SARS-CoV-2 spike proteins. PLoS Pathog. 17, e1009282 (2021).

13. P. Zhao et al., Virus-Receptor Interactions of Glycosylated SARS-CoV-2 Spike and Human ACE2 Receptor. Cell Host Microbe. 28, 586–601.e6 (2020).

14. Q. Li et al., The Impact of Mutations in SARS-CoV-2 Spike on Viral Infectivity and Antigenicity. Cell. 182, 1284–1294.e9 (2020).

15. O. C. Grant, D. Montgomery, K. Ito, R. J. Woods, Analysis of the SARS-CoV-2 spike protein glycan shield reveals implications for immune recognition. Scientific Reports. 10, 14991 (2020).

16. M. Sikora et al., Computational epitope map of SARS-CoV-2 spike protein. PLoS Comput Biol. 17, e1008790 EP – (2021).

17. E. C. Clarke, R. A. Nofchissey, C. Ye, S. B. Bradfute, The iminosugars celgosivir, castanospermine and UV-4 inhibit SARS-CoV-2 replication. Glycobiology (2020), doi:10.1093/glycob/cwaa091.

18. M. Holwerda, P. V’kovski, M. Wider, V. Thiel, R. Dijkman, Identification of five antiviral compounds from the Pandemic Response Box targeting SARS-CoV-2. bioRxiv, 2020.05.17.100404–25 (2020).

19. S. Rajasekharan et al., Inhibitors of Protein Glycosylation Are Active against the Coronavirus Severe Acute Respiratory Syndrome Coronavirus SARS-CoV-2. Viruses. 13 (2021), doi:10.3390/v13050808.

20. Q. Yang et al., Inhibition of SARS-CoV-2 viral entry upon blocking N- and O-glycan elaboration. Elife. 9, 450 (2020).

21. M. E. Dieterle et al., A replication-competent vesicular stomatitis virus for studies of SARS-CoV-2 spike-mediated cell entry and its inhibition. Cell Host Microbe (2020), doi:10.1016/j.chom.2020.06.020.

22. L. J. Scott, C. M. Spencer, Miglitol: a review of its therapeutic potential in type 2 diabetes mellitus. Drugs. 59, 521–549 (2000).

23. T. M. Cox et al., The role of the iminosugar N-butyldeoxynojirimycin (miglustat) in the management of type I (non-neuronopathic) Gaucher disease: a position statement. J Inherit Metab Dis. 26, 513–526 (2003).

24. D. Durantel, Celgosivir, an alpha-glucosidase I inhibitor for the potential treatment of HCV infection. Curr Opin Investig Drugs. 10, 860–870 (2009).

25. L. K. Campbell, J. R. White, R. K. Campbell, Acarbose: its role in the treatment of diabetes mellitus. Ann Pharmacother. 30, 1255–1262 (1996).

26. A. Helenius, M. Aebi, Roles of N-linked glycans in the endoplasmic reticulum. Annu Rev Biochem. 73, 1019–1049 (2004).

27. Ł. F. Sobala et al., Structure of human endo-α-1,2-mannosidase (MANEA), an antiviral host-glycosylation target. Proc Natl Acad Sci USA. 117, 29595 (2020).

28. J. D. Allen, Y. Watanabe, H. Chawla, M. L. Newby, M. Crispin, Subtle influence of ACE2 glycan processing on SARS-CoV-2 recognition. Journal of Molecular Biology, 166762 (2020).

29. L. C. N. da Silva et al., Exploring lectin-glycan interactions to combat COVID-19: lessons acquired from other enveloped viruses. Glycobiology (2020), doi:10.1093/glycob/cwaa099.

30. D. Hoffmann et al., Identification of lectin receptors for conserved SARS-CoV-2 glycosylation sites. bioRxiv, 2021.04.01.438087 (2021).

31. E. I. Patterson et al., Methods of Inactivation of SARS-CoV-2 for Downstream Biological Assays. J Infect Dis. 222, 1462–1467 (2020).

32. M. Kotsias et al., Method comparison for N-glycan profiling: Towards the standardization of glycoanalytical technologies for cell line analysis. PLoS ONE. 14, e0223270 (2019).

33. R. P. Kozak, C. B. Tortosa, D. L. Fernandes, D. I. R. Spencer, Comparison of procainamide and 2-aminobenzamide labeling for profiling and identification of glycans by liquid chromatography with fluorescence detection coupled to electrospray ionization-mass spectrometry. Anal Biochem. 486, 38–40 (2015).

34. L. Royle et al., HPLC-based analysis of serum N-glycans on a 96-well plate platform with dedicated database software. Anal Biochem. 376, 1–12 (2008).

35. D. Mellacheruvu et al., The CRAPome: a contaminant repository for affinity purification-mass spectrometry data. Nat Methods. 10, 730–736 (2013).

36. T. H. Plummer, J. H. Elder, S. Alexander, A. W. Phelan, A. L. Tarentino, Demonstration of peptide:N-glycosidase F activity in endo-beta-N-acetylglucosaminidase F preparations. J. Biol. Chem. 259, 10700–10704 (1984).

37. F. Maley, R. B. Trimble, A. L. Tarentino, T. H. Plummer, Characterization of glycoproteins and their associated oligosaccharides through the use of endoglycosidases. Anal Biochem. 180, 195–204 (1989).

